# An explorable public transcriptomics compendium for eukaryotic microalgae

**DOI:** 10.1101/403063

**Authors:** Justin Ashworth, Peter J. Ralph

## Abstract

Eukaryotic microalgae dominate primary photosynthetic productivity in fluctuating nutrient-rich environments, including coastal, estuarine and polar regions, where competition and complexity are presumably adaptive and dynamic traits. Numerous genomes and transcriptomes of these species have been carefully sequenced, providing an unprecedented view into the vast genetic repertoires and the diverse transcriptional programs operating inside these organisms. Here we collected, re-mapped, quantified and clustered publicly available transcriptome data for ten different eukaryotic microalgae in order to develop new insights into their molecular systems biology, as well as to provide a large new resource of integrated information to facilitate the efforts of others to further compare and contextualize the results of individual and new experiments within and between species. This is summarized herein and provided for public use by the eukaryotic microalgae research community.

## Introduction

Eukaryotic microalgae dominate primary photosynthetic productivity in fluctuating nutrient-rich environments, including coastal, estuarine and polar regions, where competition and complexity are presumably adaptive and dynamic traits [1–3]. Numerous genomes and transcriptomes of these species have been carefully sequenced, providing an unprecedented view into the vast genetic repertoires and the diverse transcriptional programs operating inside these organisms [4–16]. The integration of these transcriptome data to allow wholistic, multi-experiment and system-level analysis and interpretation is important and fruitful for gaining new understandings of concerted microbial functions [17,18].

In order to facilitate new insights into the molecular systems biology of eukaryotic microalgae, we systematically collected, re-mapped, quantified and clustered publicly available transcriptome data for ten different eukaryotic microalgal species, investigated methods and caveats of integration, performed basic clustering, functional annotation and orthology analyses within and between species, and built an interactive online resource for the further exploration of the largest single compendium of microalgal transcriptomics data to date.

## Results

### Data integration

The sequencing read data for nine microalgal species were obtained from the Sequence Read Archive (SRA) [19] and systematically re-mapped to current genome assemblies using HISAT2 [20], SAMtools [21], and StringTie [22] on Amazon Web Services [23] (see Materials and Methods). By-sample transcriptomic read counts for *Thalassiosira weissflogii* were estimated by mapping reads onto genome-free transcript assemblies from the Marine Microbial Eukaryote Sequencing Project [9]. Transcriptomic data for the freshwater microalga *Chlamydomonas reinhardtii* are the subject of ongoing study by other groups [24], and may be combined with these data in the future. Within-species normalization of raw transcript-per-million (TPM) [25] counts was conducted using a method developed to maximize the number of uniform genes [26].

### Batch effects

The integration of independent transcriptome experiments is prone to systematic variations that are unique‐‐but often consistent‐‐to each individual laboratory, treatment, preparation, platform, or other unaccountable factors. It is presumed that some of these biases may be corrected by applying an appropriate normalization to the data, while others may remain xs[27].

To normalize RNA-seq transcriptome data, we applied a method developed to algorithmically maximize the number of “uniform genes,” or those for which transcript levels are least variant over multiple experiments, to serve as internal standards for within-sample normalization factors [26]; this compared favourably in this regard to the ‘Trimmed Mean of M Values’ (TMM) [28] and other methods of normalization across a large RNA-seq dataset of diverse human tissues. Application to microalgal transcriptome data in this study adjusted distributions of log_10_TPM values across different experimental series (Supplementary Figure S1), resulting in improvements to the consistency of measurements for typical “housekeeping” genes, such as actins, glyceraldehyde 3-phosphate dehydrogenase (GAPDH), tubulins, and ribosomal proteins (Supplementary Figure S2). Remaining batch effects apparent between different studies included sets of transcripts whose within-study biases may be attributable to biological and/or non-biological sources.

Previously assembled microarray data for *Thalassiosira pseudonana* and *Phaeodactylum tricornutum* [29] were appended to this dataset to facilitate direct comparisons between the results of different platforms used to estimate transcriptome-wide expression levels, as well as between independent experiments. Microarray fold-change ratio data and RNA-seq TPM data in aggregate were distributed similarly (Supplementary Figure S3). These data are presented and provided without further normalisation to fit ideal distributions, and therefore preserve biologically faithful measurements and to facilitate further comparisons between other similarity, normalisation and clustering approaches.

In total, 1,375 samples (transcriptome measurements) were integrated from sixty-nine independent experimental studies representing ten species and four distinct clades of microalgae (Figure 1, Supplementary Table S1), providing a rich consolidated dataset for new and comparative data exploration.

**Figure 1.**
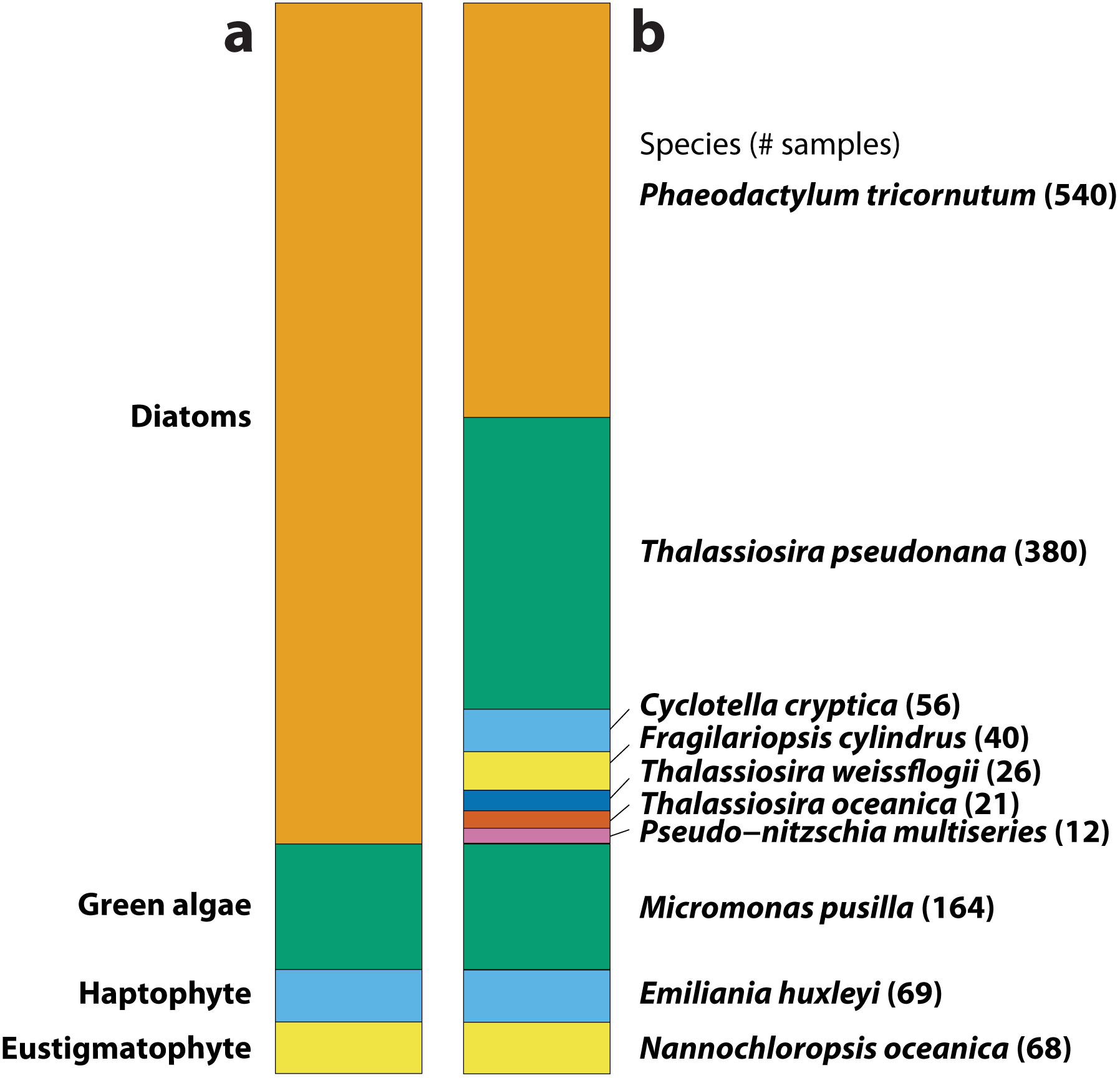
Summary of sample counts by a) microalgal clade and b) species.

### Clustering with efficient empirical estimates of reproducibility

Agglomerative hierarchical clustering based on aggregate within-sample correlations [30] was performed for each species, resulting in high numbers of putative clusters with distinct conditional and experimental patterns of expression in all ten microalgal species. In order to assess the robustness, reproducibility and statistical boundaries of apparent clusters of coexpressed transcripts, we employed deep bootstrapping of hierarchical trees with multi-scale resampling using a version of Pvclust [31] refactored to run more efficiently on large matrices [29]. This produced data-supported, fine-grained clusters of transcripts (Supplementary Table S1) whose expression patterns over all conditions imply‐‐but not do prove‐‐uniquely shared responses, regulation, activities, or functions that appear to be linked to biological or environmental change. Large, robust clusters of co-expressed and putatively related genes include major aspects of microalgal biology, such as the regulation of light harvesting and ribosomes, while smaller clusters of condition-specific genes abound, such as those for nutrient scavenging (Figure 2).

**Figure 2.**
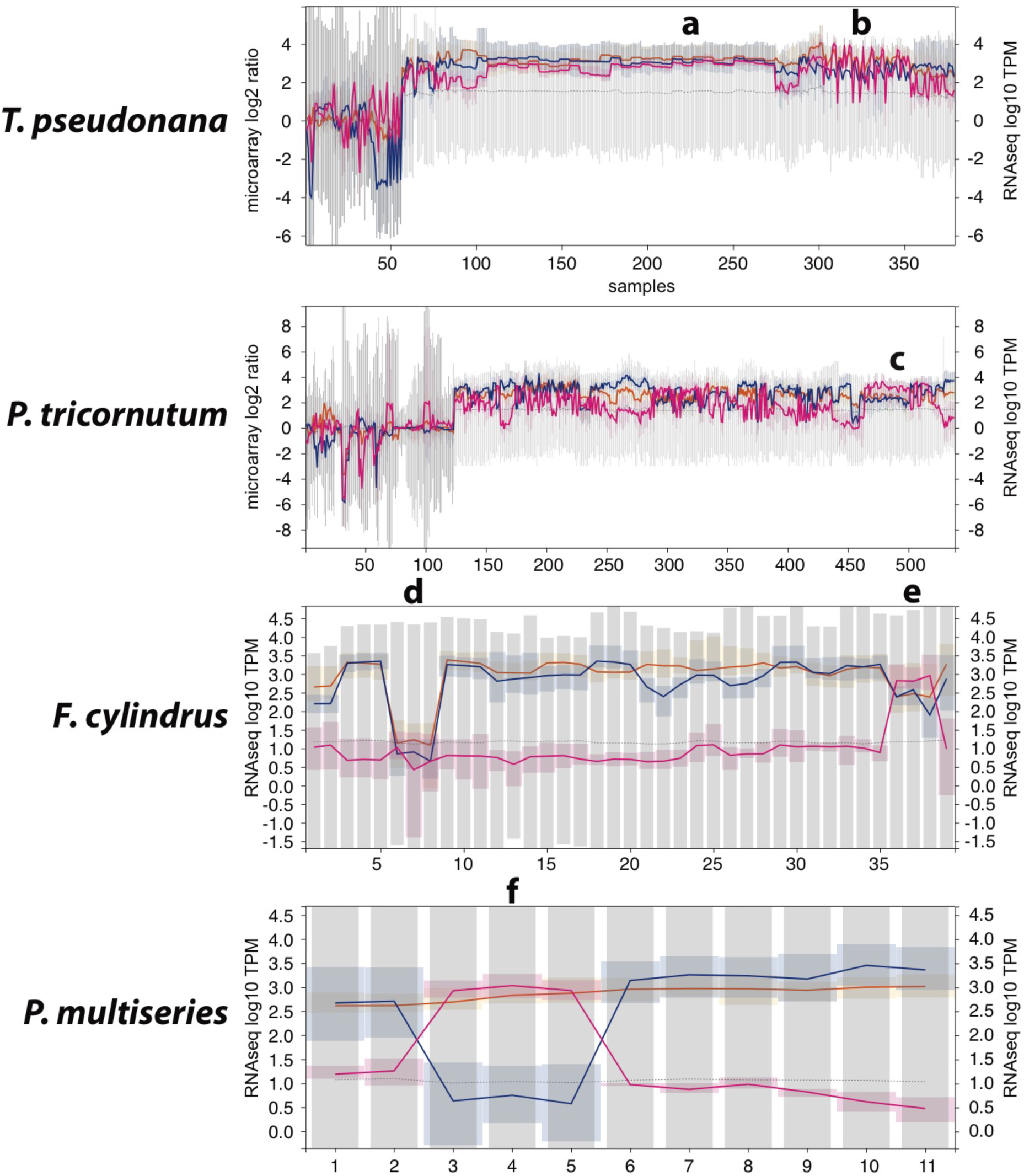
Distinct bootstrap-supported clusters of co-expressed genes in four diatom species, which include ribosomal proteins (orange), photosynthesis and light harvesting genes (blue), and nitrate uptake genes (magenta). Notable conditions affecting the transcript levels of these gene groups include a) nitrate-limited chemostat cultures of *T. pseudonana* [32], b) batch cultures of *T. pseudonana* experiencing periodic nutrient stress [33], c) *P. tricornutum* cultures subject to nitrate depletion [34], *F. cylindrus* subject to d) prolonged darkness [12] and e) nitrate limitation [35], and f) *Pseudo-nitzschia multiseries* subject to nitrate limitation [35].

The occurrence of orthologically related clusters of co-expressed genes across all included species reflects a conservation of concerted transcriptional regulation of fundamental microalgal processes. In the case of nitrate uptake mechanisms in diatoms (Figure 2, magenta lines), the transcript levels of putatively orthologous nitrate uptake transporters correlate with organic nitrogen-limited conditions in multiple species. An abundance of smaller apparent clusters, comprised of large proportions of novel and under-characterized genes, implies a wealth of divergently regulated functions whose unique expression patterns might contribute to species specificity and environmentally linked biological programs.

### Public exploration via online resource

To further increase the public utility and accessibility of this data resource, we created a lightweight database that includes basic protein homolog predictions using BLASTp [36], protein sequences, functional annotations, putative promoter region sequences, as well as an interactive online interface to this compendium using Django [37] and d3.js [38]. This resource (Figure 3) is currently available at https://alganaut.uts.edu.au in cooperation with the Australian National eResearch Collaboration Tools (NeCTAR) [39]. The transcriptomic datasets assembled in this work are also available for download, independent analysis and use.

**Figure 3.**
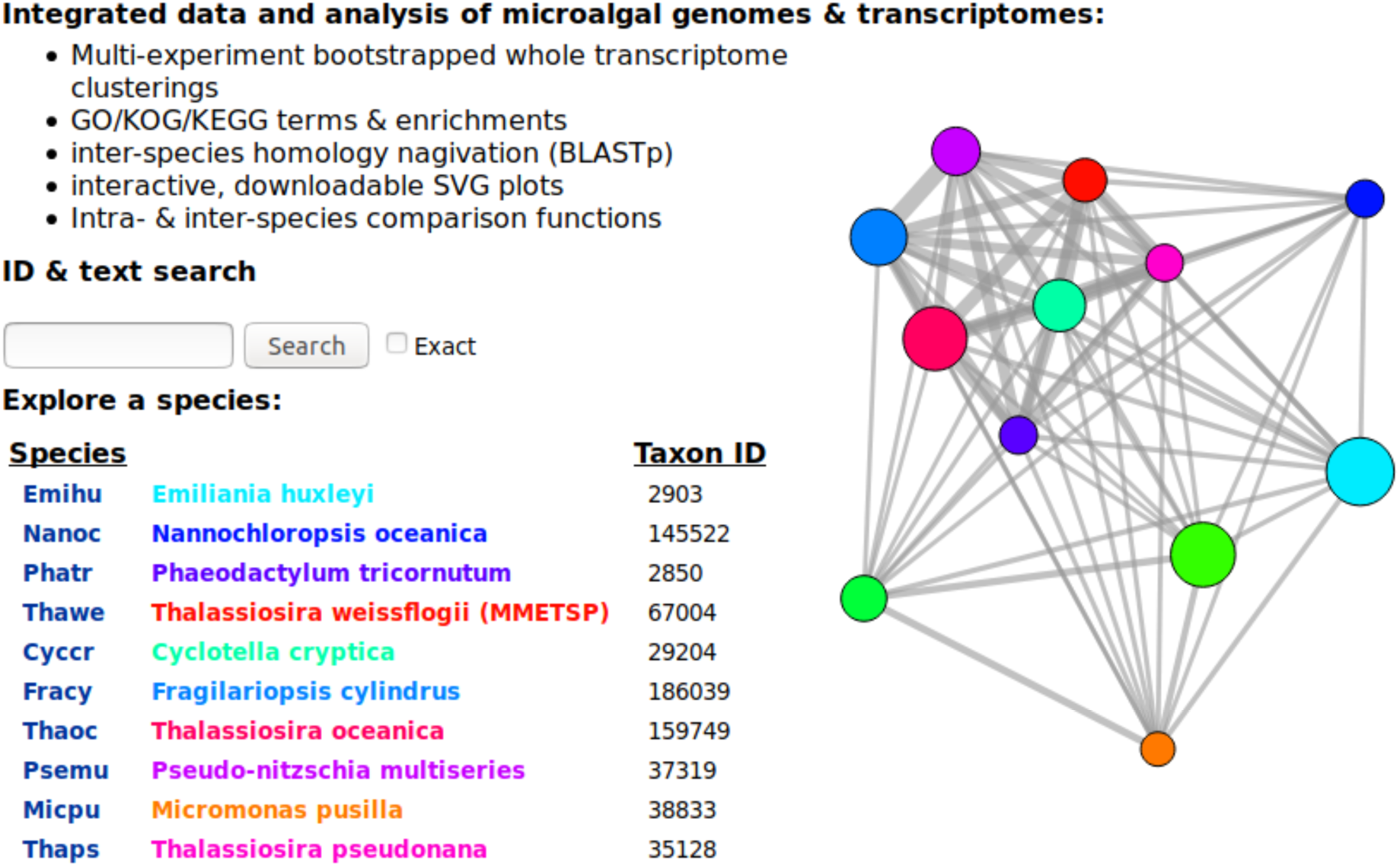
The “Alganaut” public interactive web resource for the exploration of microalgal transcriptomes.

## Discussion

Assembly and informed analysis of ever-expanding integrated datasets are crucial for organismal and environmental sciences. While dozens of microalgal genomes have now been sequenced, assembled and translated into large and diverse proteomes, the identities and functions of the majority of putative proteins discovered in these organisms remain a mystery. Approximately half or more of each new microalgal proteome bears no reliable similarity to any other species, nor to any known proteins or functional domains in existing gene databases [4–13]. Furthermore, mechanistically accurate gene networks, molecular “biomarkers,” robust environmental covariates, and intra‐ and inter-species dependencies are all yet to be discovered for eukaryotic microalgal species. “Data-driven” large-scale prediction and discovery of putative links between known and unidentified proteins will assist in further modeling, characterization and understanding of algal species.

### Data science potential

In most species, the transcript levels of nearly all functional gene products depend on the physiological state of the cell or cell population, the biological and regulatory programs in operation, and environmental conditions. It is presumed that groups of transcripts that exhibit uniquely similar or correlated expression patterns over various conditions are more likely to be functionally related or co-regulated than those for which expression patterns diverge.

Mining of transcriptome data can be performed in numerous ways. Most approaches yield a finite number of large clusters with similar memberships based on various algorithms, as well as increasingly divergent sub-clusters based on various arbitrary, heuristic, or statistical choices. Additional types of clustering and significance estimation can be fruitfully applied to these data. Both a) empirically appropriate statistical metrics and b) new and orthogonal experimental measurements will be required to agnostically compare the results of different “data-driven” predictions, as has been accomplished for established bacterial, metazoan and human systems [40]. Sources of validation and extension of models include fresh transcriptional data series, gene knockdowns, knockouts, overexpression, genome-wide binding studies, comparative genomics, proteomics, and subcellular localization atlases. Increasing amounts of these orthogonal experimental measurements will be particularly valuable to provide a sufficient scientific basis upon which to judge the accuracy and applicability of biological and statistical models.

### Limitations of transcriptome data

Transcriptomics alone is insufficient to understand the complex biology and inner workings of microalgae. However, it is currently the easiest approach to gather comprehensive data to describe the intracellular behaviour, cellular programming, environmental responses, and comparative molecular biology. The ease of obtaining, handling and interpreting sequence read data continues to outpace other types of measurements, including proteomics, phenomics, fluorometry, cytometry, and imaging. But the value of genomic and transcriptomic data are limited without adequate context‐‐thus the relative shortage of other data types such as those mentioned above make them increasingly valuable to collect in parallel with transcript data.

Post-transcriptional regulation and post-translational features including RNA modifications, binding and trafficking, degradation, protein modifications, subcellular localization, protein and chemical signalling, and allostery all crucially contribute to the cellular holo-program. Protein levels do, as an apparent rule, correlate closely with their corresponding transcript levels [42,43], but it is possible to identify interesting exceptions. For example, peptide signalling is prevalent in eukaryotic microalgae [44,45], and the complex subcellular locations of variously acting proteins is crucial to correctly understand their physiological roles across a large number of different membranes and compartments [46]. The measurement of protein phospho-states [47,48], which can provide data to model protein signalling and the post-translational activities of many transcription factors, may also be possible and relevant to advance systems biology studies in microalgae.

### Integrated datasets for environmental biology

Biology is governed by detailed regulatory and metabolic programs operating throughout various environmental conditions. While rapid, high-throughput reverse genetics and phenotyping approaches are under development eukaryotic microalgal species, multi-omics and experimental systems biology are the fastest and broadest initial means to observe and predict the roles, functions and importance of new genes in new and non-model organisms. Molecular systems biology consists of collecting detailed and comprehensive measurements of numerous observable molecular features and parameters, and then using computing, statistics, quantitative hypotheses and biological models to apply these data to new questions [49]. The synthesis of community-wide datasets will continue to deepen our understanding of microalgae and help to inform further efforts including biological oceanography, direct genetics, and comparative microbial biology, and biotechnology.

## Materials and Methods

### RNA-seq data integration and normalization

Sequencing read data for eukaryotic microalgae with sufficient numbers of samples for integration (≥10, Supplementary Table S2) were obtained from the Sequence Read Archive (SRA) [19] as of February 2018 and re-mapped to current genome assemblies using HISAT2 v2.1.0 [20], SAMtools 1.7 [21], and StringTie 1.3.4b [22] on Amazon Web Services [23]. Genome and RNA-seq data for *C. cryptica* were obtained from Traller et. al [11]. By-sample transcriptomic read counts for *T. weissflogii* were estimated by mapping reads onto genome-free transcript assemblies from the Marine Microbial Eukaryote Sequencing Project [9]. The ordering and sample names of data series were taken as-is in accordance with source annotations. Within-species normalization of raw transcript-per-million (TPM) [25] counts was conducted using a method developed to maximize the number of uniform genes [26]. Previously assembled microarray data for *T. pseudonana* and *P. tricornutum* [29] were appended as log_2_ ratios of changes in expression to within-experiment internal reference controls. Various scripts used and intermediate datasets are available as Supplementary Information.

### Bootstrapped hierarchical clustering

Agglomerative hierarchical clustering was performed using a c++ wrapper to call the Fastcluster [30] library directly and repeatedly, with in-memory multi-scale resampling of the data to perform efficient bootstrapping [29]. A partially refactored version of Pvclust [31] was used to assign bootstrap P-values to branch nodes in the hierarchical tree, providing a robust empirical estimate of reproducible and bifurcating sub-cluster memberships.

### Database and resource implementation

An SQL database was used to combine transcriptomic data with pairwise reciprocal inter-proteome homolog predictions (BLASTp 2.6.0 [36]), GO, KEGG, and KOG functional annotations where publicly available from draft genomes, and putative upstream promoter region sequences taken from public genome assemblies. These data were rendered explorable using Django 2.0.3 [37] and d3.js version 4 [38]. The data resource was made available on-line using a remote server instance on the Australian NeCTAR Research Cloud [39].

## Acknowledgements

J.A. is the recipient of an Australian Research Council Discovery Early Career Award (DE160100615) funded by the Australian Government. This research was supported by use of the NeCTAR Research Cloud, a collaborative Australian research platform supported by the National Collaborative Research Infrastructure Strategy.

---

**Table S1.**
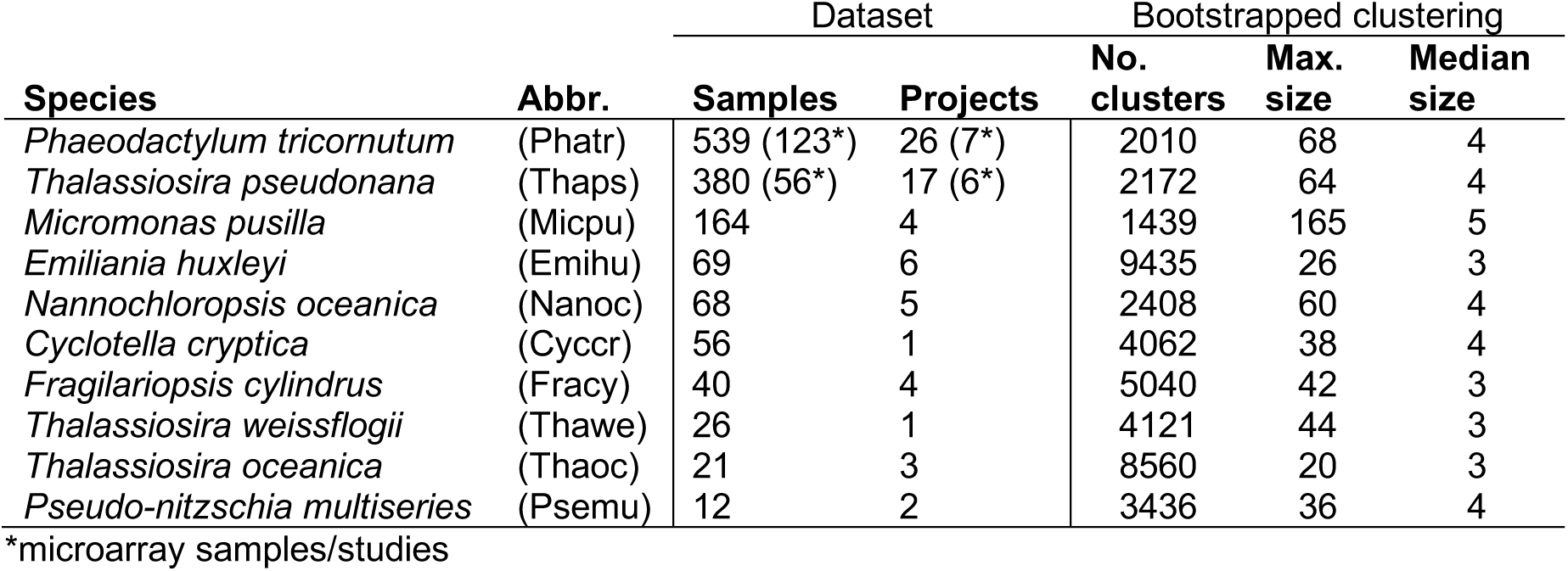
Dataset summary.

**Table S2.** RNA-seq samples included in dataset (separate file).

**Figure S1.**
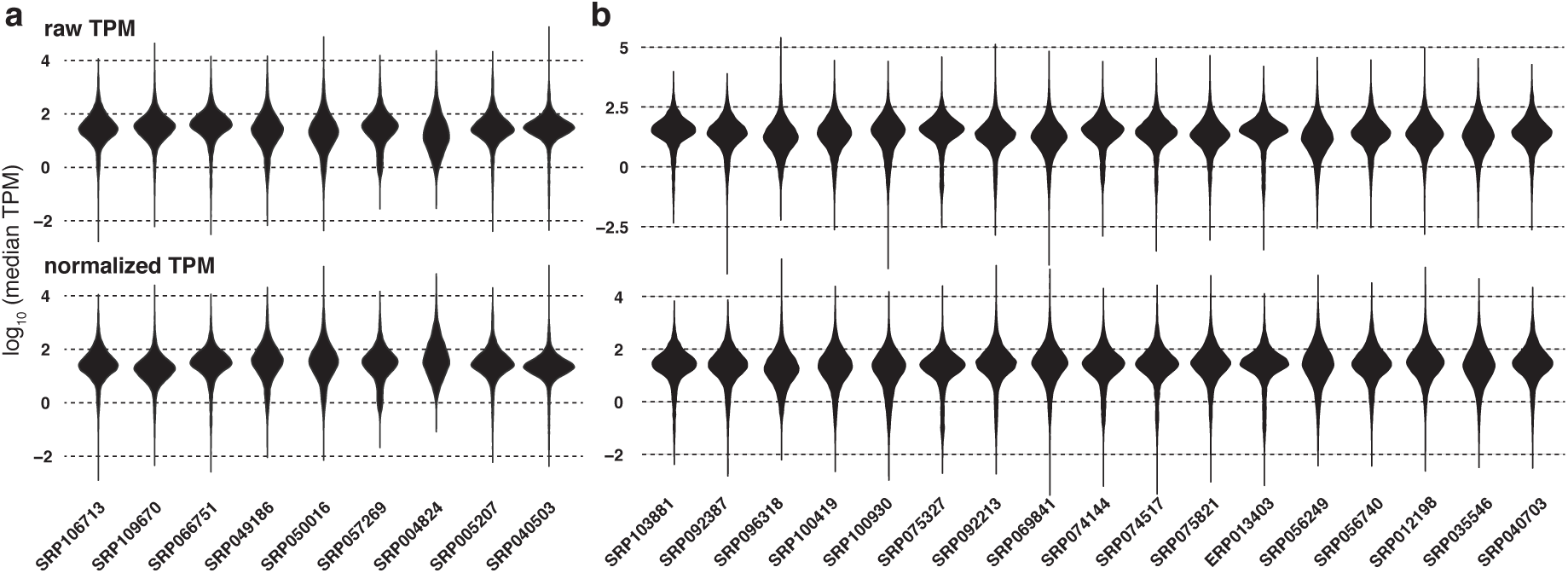
Log_10_ by-transcript median TPM distributions (top: raw, bottom: normalized) for different RNA-seq experimental project series, for a) *T. pseudonana* and b) *P. tricornutum*.

**Figure S2.**
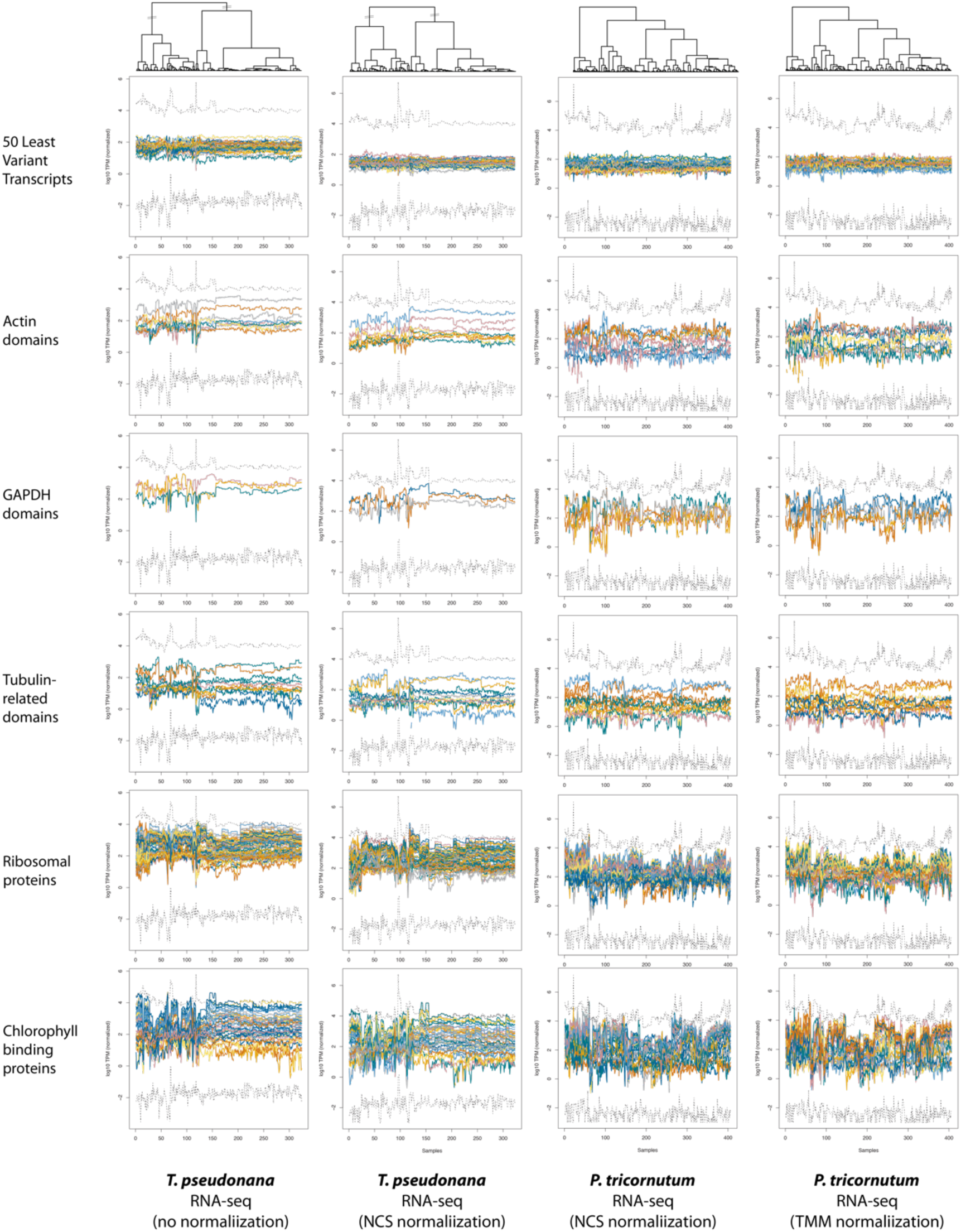
Combined RNA-seq datasets for *T. pseudonana* and *P. tricornutum*, illustrating characteristic results of two routine batch normalization algorithms: “network centrality scaling” (NCS) and “trimmed mean of means” (TMM). The transcripts per million (TPM, yaxis) for individual transcripts over all samples (x-axis) are shown as arbitrarily colored lines. Dashed lines indicate the within-sample minimum and maximum range of TPM values over all transcripts.

**Figure S3.**
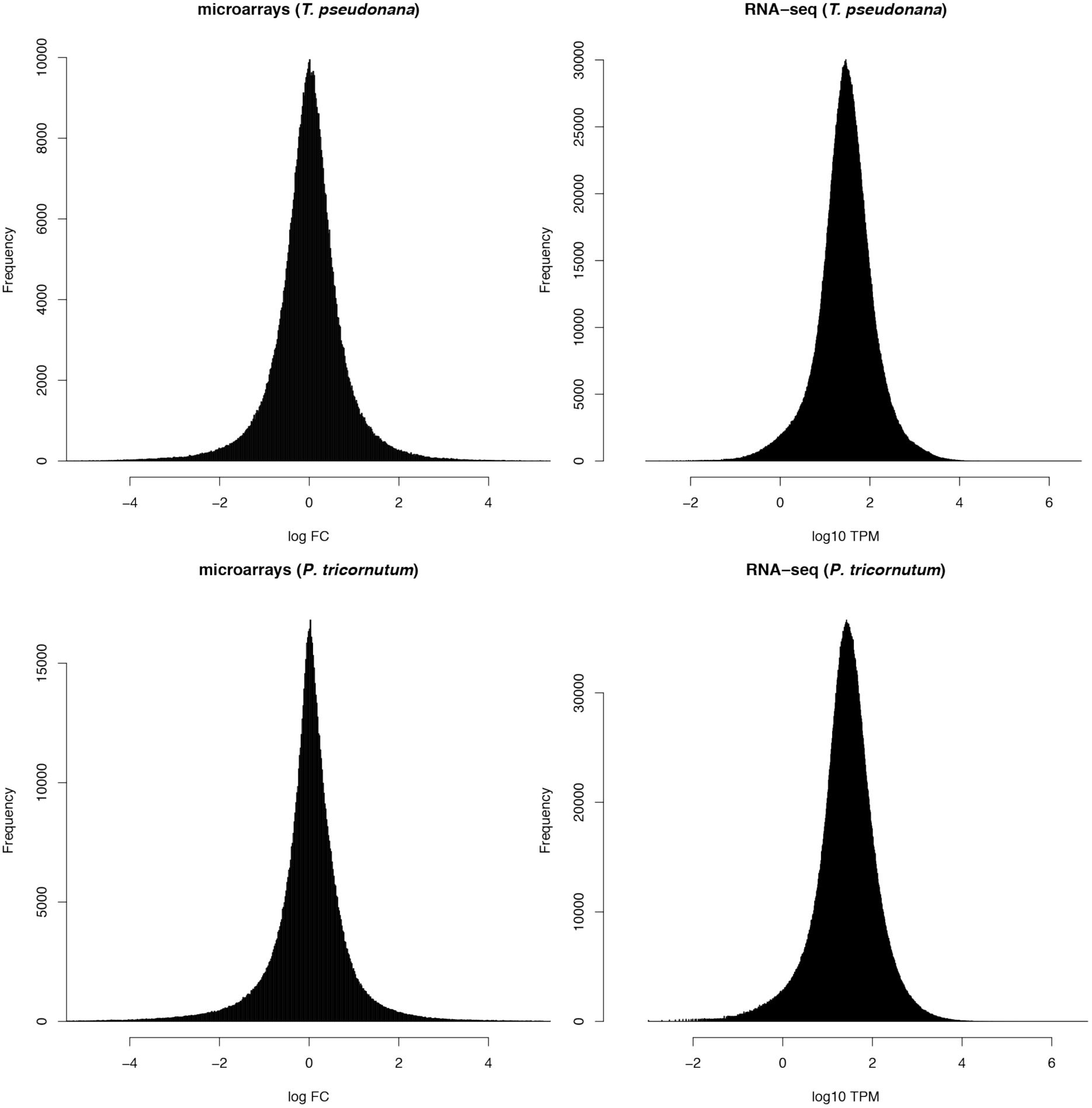
Distributions of microarray fold change (FC) (left) and RNA-seq normalized transcripts per million (TPM) values (right) included in the integrated datasets for *T. pseudonana* (top) and *P. tricornutum* (bottom).

